# Identifying long non-coding RNAs and characterizing their functional roles in swine mammary gland from colostrogenesis to lactogenesis

**DOI:** 10.1101/2021.01.17.426976

**Authors:** Lijun Shi, Longchao Zhang, Ligang Wang, Xin Liu, Hongmei Gao, Xinhua Hou, Hua Yan, Fuping Zhao, Wentao Cai, Lixian Wang

**Affiliations:** Institute of Animal Science, Chinese Academy of Agricultural Sciences, Beijing 100193, China

**Keywords:** Swine, Mammary gland, Colostrum, LncRNA, WGCNA

## Abstract

Long non-coding RNAs (lncRNAs) play important regulatory roles in milk biological processes. While, little is known about the identification and function of swine lncRNAs in response to mammary gland development. Here, we identified 286 differentially expressed (DE) lncRNAs in mammary gland at different stages from 14 days prior to (-) parturition to day 1 after (+) parturition using the published RNA-seq data. The expression pattern of these DE lncRNAs was examined, and most of lncRNAs expressions were strongly changed from Day −2 to Day +1. Functional annotation revealed that the DE lncRNAs were mainly involved in the mammary gland developing, lactation developing, milk composition metabolism and colostrum function. By performing the weighted gene co-expression network analysis (WGCNA), we identified 7 out of 12 lncRNA-mRNA modules, including 35 lncRNAs and 319 mRNAs, which were highly associated with the mammary gland at Day −14, Day −2 and Day +1. Integrated analysis of the DE lncRNAs expression pattern examination, targets prediction, function annotation and WGCNA suggested that 18 lncRNAs (such as, *XLOC_020627, ENSSSCG00000051193, XLOC_025150, ENSSSCG00000042618, XLOC_963181, ENSSSCG00000051701, XLOC_018030*, and *XLOC_025146*) and their 20 target genes (such as, *ACTN4, ADCY1, CSN3, SMO, CSN2, PRKAG2, FIBCD1*, and *GALNT7*), were considered as the promising candidates for swine parturition and colostrum occurrence processes. Our research provided the insights into lncRNA profiles and their regulating mechanisms from colostrogenesis to lactogenesis in swine.

## Introduction

Although the litter size of sow has made a significant improvement during last two decades, the problem of growth performance and mortality in piglets become increasingly prominent. During gestation and lactation, the growth performance and health status of piglets have been critical factors impacting reproductive performance of modern sows (Kim 1999). Sow milk performance as a major limiting factor contributes to suboptimal growth and survival of piglets (Boyd & Kensinger 1998). The quality and quantity of milk in sows are highly variable, for example, the colostrum yield is reported to range from <1kg to 8.5kg (Quesnel et al. 2012; Vadmand et al. 2015). To enhance the sow lactation ability, it is necessary to understand the mammary gland development, and screen their regulatory factors (Hurley 2019). In 2016, Balzani *et al*. reported the heritability of udder morphology in crossbreed sows (Large White × Meishan), namely, teat length with *h*^*2*^ = 0.46, teat diameter with *h*^*2*^ = 0.56, interteat distance within the same row with *h*^*2*^ = 0.37, abdominal midline with *h*^*2*^ = 0.22, udder development score *h*^*2*^ = 0.25, proportion of nonfunctional teats with *h*^*2*^ = 0.3, and the proportion of teats oriented perpendicular to the udder with *h*^*2*^ = 0.1 (Balzani et al. 2016). Generally, from colostrogenesis to lactogenesis, mammary glands might undergo significant functional differentiation for swine parturition and colostrum occurrence processes. In 2018, Palombo *et al*. used the RNA sequencing (RNA-seq) to detect the candidate genes impacting swine parturition and lactation in mammary gland of crossbred sows (Danish Landrace × Yorkshire), such as *CSN1S2, LALBA, WAP, SAA2*, and *BTN1A1* (Palombo et al. 2018).

Long non-coding RNA (LncRNA) is a newfound class of non-protein coding transcripts in eukaryotes with a minimum length of 200 nt (Jin et al. 2020), and it plays important regulatory role in biological processes (Liang et al. 2018). Many lncRNAs have been reported along with the depth and quality of RNA-seq (St Laurent et al. 2015). In swine, lots of potential regulatory lncRNAs have been identified from various tissues, such as intramuscular adipose (Huang et al. 2018; Miao et al. 2018), longissimus dorsi muscle (Wang et al. 2019a), preadipocytes (Li et al. 2019), and porcine endometrium (Wang et al. 2017). In 2018, Liang *et al*. built a systematic S.scrofa lncRNA database named lncRNAnet that contained 53,468 S.scrofa lncRNAs with their sequence characteristics, genomic locations, conservation, overlapping single nucleotide polymorphism and quantitative trait loci, and transcript abundance across nine tissues (fat, heart, kidney, liver, lung, muscle, ovarium, spleen, and testis) (Liang *et al*. 2018). However, no study has been performed to detect the lncRNAs regulating milk traits from sow mammary gland.

In the present study, we identified lncRNAs in sow mammary gland at different stages from 14 days prior to parturition to day 1 after parturition according to the published RNA-seq data, and detected the differentially expressed (DE) lncRNAs. Further, functional annotation and WGCNA were conducted to predict the regulatory functions of DE lncRNAs. To the best of our knowledge, this is the first study to identify lncRNAs from mammary gland in swine, and our results paved the way for a better understanding of lncRNA functions in swine parturition and colostrum occurrence processes.

## Material and methods

### RNA-seq dataset

The RNA-seq data sources (GSE101983) were involved in 15 mammary gland tissues, which were collected from 3 crossbred sows (Danish Landrace × Yorkshire) at Days 14, 10, 6 and 2 before (-) parturition and Day 1 after (+) parturition (Table S1)(Palombo *et al*. 2018).

### Identification of lncRNAs

According to the raw data, clean reads were obtained by removing containing adapter molecules, reads including poly-N, and low-quality reads using Trimmomatic with defaults parameters (Bolger et al. 2014). All follow-up analyses were performed with the high-quality reads. Through STAR software, we aligned the clean reads to the pig reference genome (Sscrofa11.1).

We used StringTie and Scripture to assemble the novel transcripts. All the transcripts with the same start, end position and exon–intron boundary, which were supported by at least two assembly programs or occurred in at least two samples were obtained (Guttman et al. 2010; Pertea et al. 2015). Further, the putative novel lncRNAs were detected via a customized multi-step pipeline: (1) The transcripts were removed, which were likely to be assembly artifacts or PCR run-on fragments based on class code annotated by gffcompare. For the different class, we retained those only annotated by ‘u’ and ‘i’, which indicated novel intergenic and intronic transcripts, respectively.

Transcripts with class code “=“were considered as known genes. (2) Transcripts with length ⩾ 200 nt and exon ⩾ 2 were retained to avoid incomplete assemble and too many splicing events. (3) Transcripts with low expression were removed using FPKM ⩾ 0.3 as cut off. The expression levels of transcripts were calculated in fragments per kilo-base of exon per 10^6^ mapped fragments (FPKM) by StringTie −e −B. (4) Maximum ORF (open reading frame) lengths of less than 120 amino acids (360 nt) were obtained. (5) Transcripts with predicted protein-coding potential were removed, that the protein-coding potential criteria were CPC score > 0, PLEK score > 0, and CNCI score > 0. Finally, the candidate sets of lncRNAs were made up by those without coding potentials.

### Differential expression analysis of lncRNAs

To detect the DE lncRNAs and coding genes, the expression levels of these lncRNAs and coding genes identified in this study from the mammary gland at Days − 10, −6, −2 and +1 were compared to their respective expression levels at Day −14 using the EdegR R package. RNA-Seq read counts were modeled by a generalized linear model that considered the experimental design with two factors (individuals and stages of lactation). Therefore, the formula in EdegR showed as follow: Design= ∼Individuals + Stages. LncRNAs or genes with adjusted *P*-value < 0.05 were assigned as the differences. The expression patterns of DE lncRNAs across five stages were performed by k-mean method (Ahmad & Dey 2007). Through gap statistics, we determined that k = 14 was the optimal choice for distinguishing these lncRNAs.

### Target gene prediction of lncRNAs and functional analysis

LncRNAs can cis-regulate neighboring target genes and trans-regulate distant target genes (Derrien et al. 2012; Guil & Esteller 2012). To predict cis-regulated target genes of lncRNAs, the coding genes located within 100 kb upstream/downstream of lncRNAs were checked. Further, we computed the expressed correlation coefficients between lncRNAs and their neighboring genes by Spearman method, in which, the significant lncRNA-mRNA correlation pairs were assigned when the *P*-value < 0.05.

To investigate the functions of DE lncRNAs, we performed gene ontology (GO) and kyoto encyclopedia of genes and genomes (KEGG) enrichments using KOBAS (http://kobas.cbi.pku.edu.cn/kobas3/genelist/) (Xie et al. 2011). The significant GO terms and pathways were assigned with the *P*-value < 0.05.

### Module construction base on WGCNA

WGCNA is a systematic in silico method for analysis of complex gene regulatory networks, which can be used for finding modules of significantly correlated genes, and for relating modules to one another and to external sample traits (Langfelder & Horvath 2008; Le et al. 2019). Here, we conducted WGCNA to detect the significant modules by WGCNA R package. In the significant modules, we screened the lncRNAs and genes, which might be as candidates to impact the swine lactation developing, milk fat and protein metabolism and colostrum function.

## Results

### Identification of lncRNAs

In this study, RNA-seq data of 15 samples from the mammary gland of three crossbred sows with five time points were used (Table S1) (Palombo *et al*. 2018). After quality control, reads mapping and lncRNA identification, a total of 1,346 lncRNA transcripts located in 1,084 lncRNA loci were detected (Table S2), in which, 483 (613 lncRNA transcripts) were novel lncRNAs, and 601 (733 lncRNA transcripts) were known lncRNAs.

### Differential expression analysis

A total of 286 lncRNAs (Table S3) were differentially expressed (adjusted *P*-value < 0.05), including 9 (7 up-regulated and 2 down-regulated), 71 (41 up-regulated and 30 down-regulated), 169 (101 up-regulated and 68 down-regulated) and 206 (128 up-regulated and 78 down-regulated) for −10vs-14, −6vs-14, −2vs-14 and 1vs-14 comparison groups, respectively (Figure 1A-D). Notably, six lncRNAs were differentially expressed in all four comparison groups (Figure 2). Further, we examined the expression pattern of DE lncRNAs across five lactation stage using k-means clustering, and these lncRNAs could be divided into 14 distinct clusters. In cluster 2, 3, 7, 12, 13 and 14, most of DE lncRNAs expressions were rapidly decreased at Day −2, and strongly increased at Day +1, which indicated these lncRNAs were dynamic changed from colostrogenesis to lactogenesis (Figuer 3).

**Figure 1.**
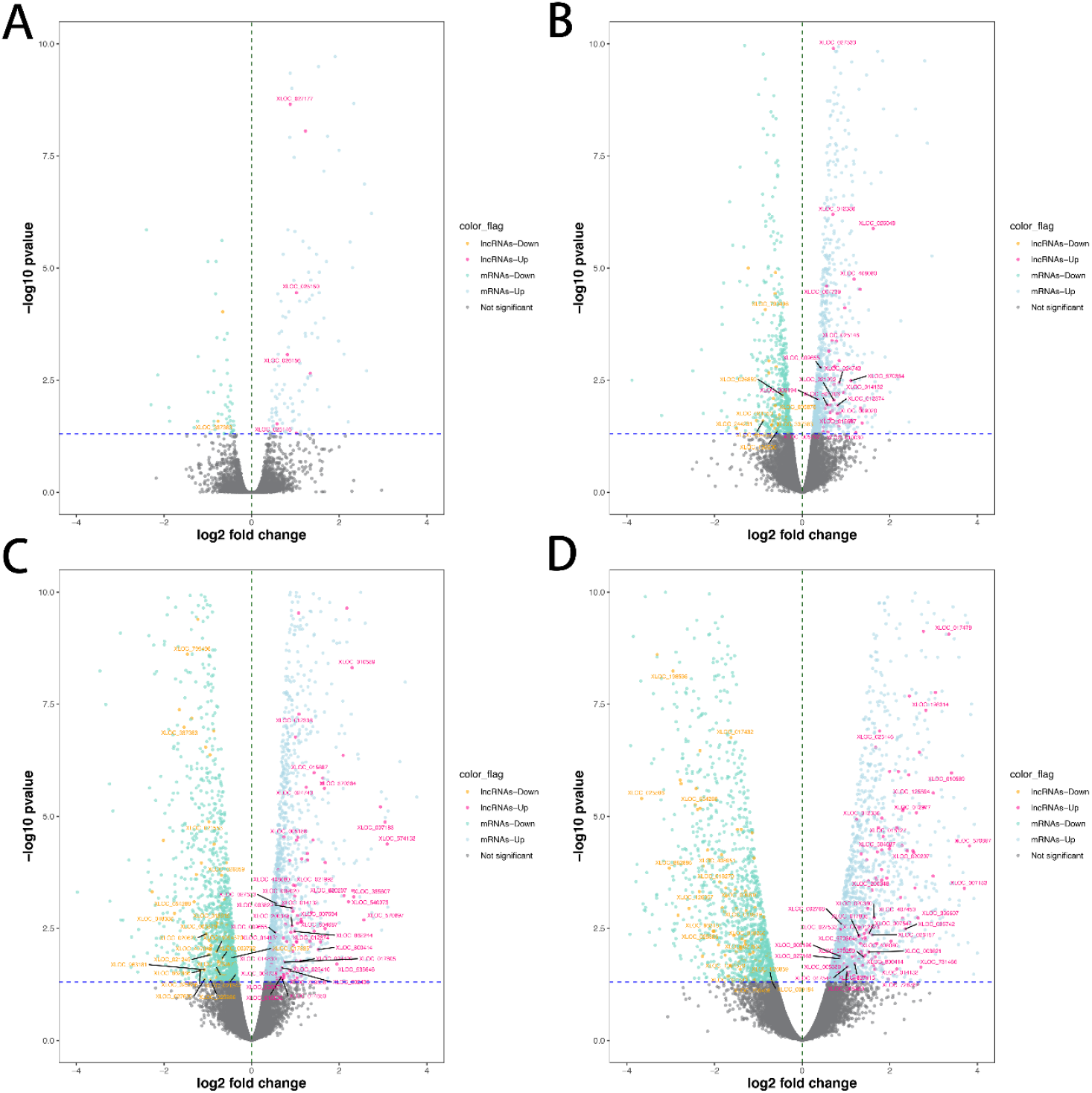
Identification of differentially expressed (DE) lncRNAs and mRNAs. **(A)** Volcano plot dispalys DE lncRNAs and mRNAs of Day −10 vs Day −14. **(B)** Volcano plot dispalys DE lncRNAs and mRNAs of Day −6 vs Day −14. **(C)** Volcano plot dispalys DE lncRNAs and mRNAs of Day −2 vs Day −14. **(D)** Volcano plot dispalys DE lncRNAs and mRNAs of Day +1 vs Day −14. The novel DE lncRNAs was marked.

**Figure 2.**
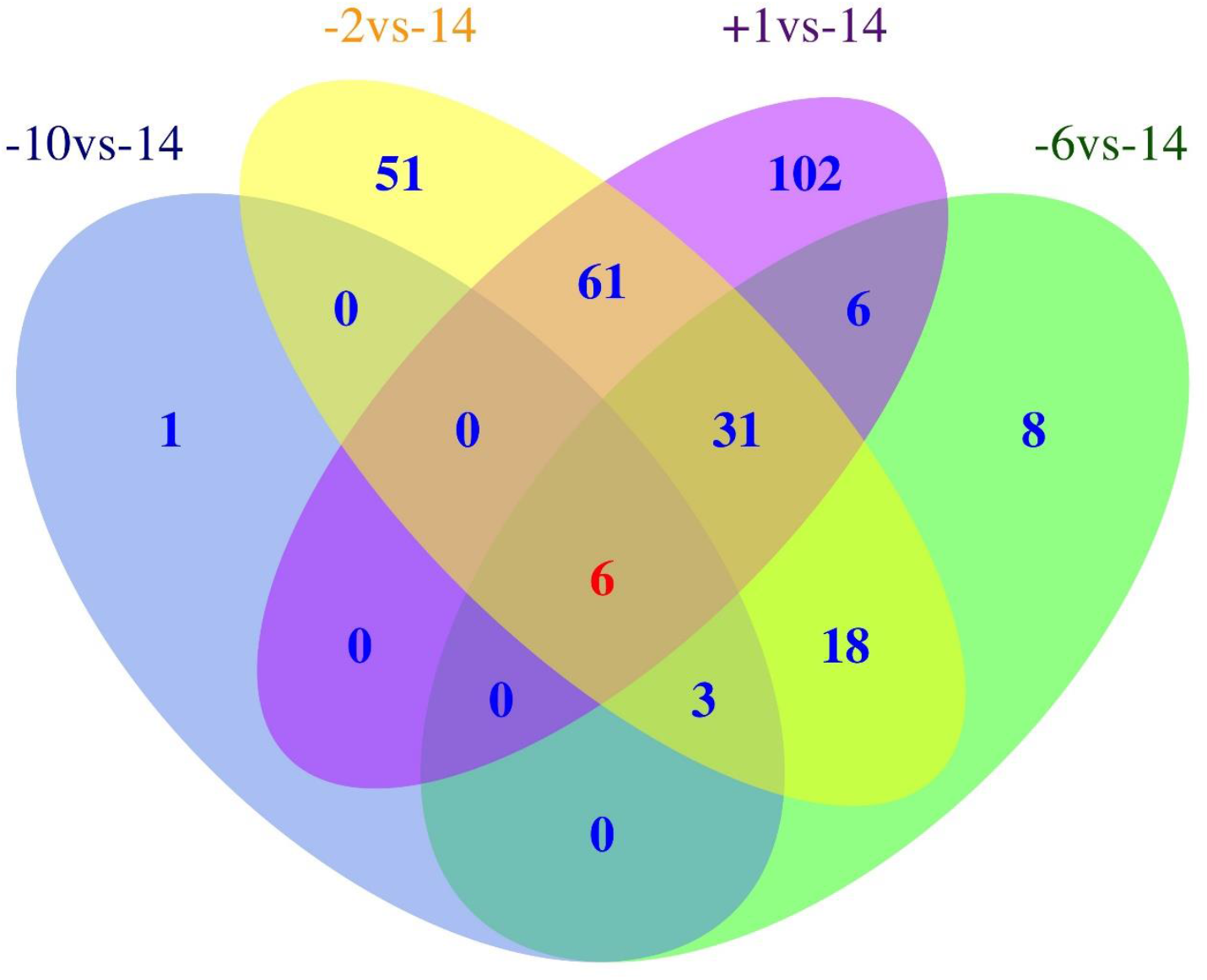
Venn diagram shows the number of differentially expreesed (DE) lncRNAs overlapping in different swine mammary gland development stages or unique to each developmental stage. -14: at days 14 before parturition. −10: at days 10 before parturition. −6: at days 6 before parturition. −2: at days 2 before parturition. +1: at days 1 after parturition.

**Figure 3.**
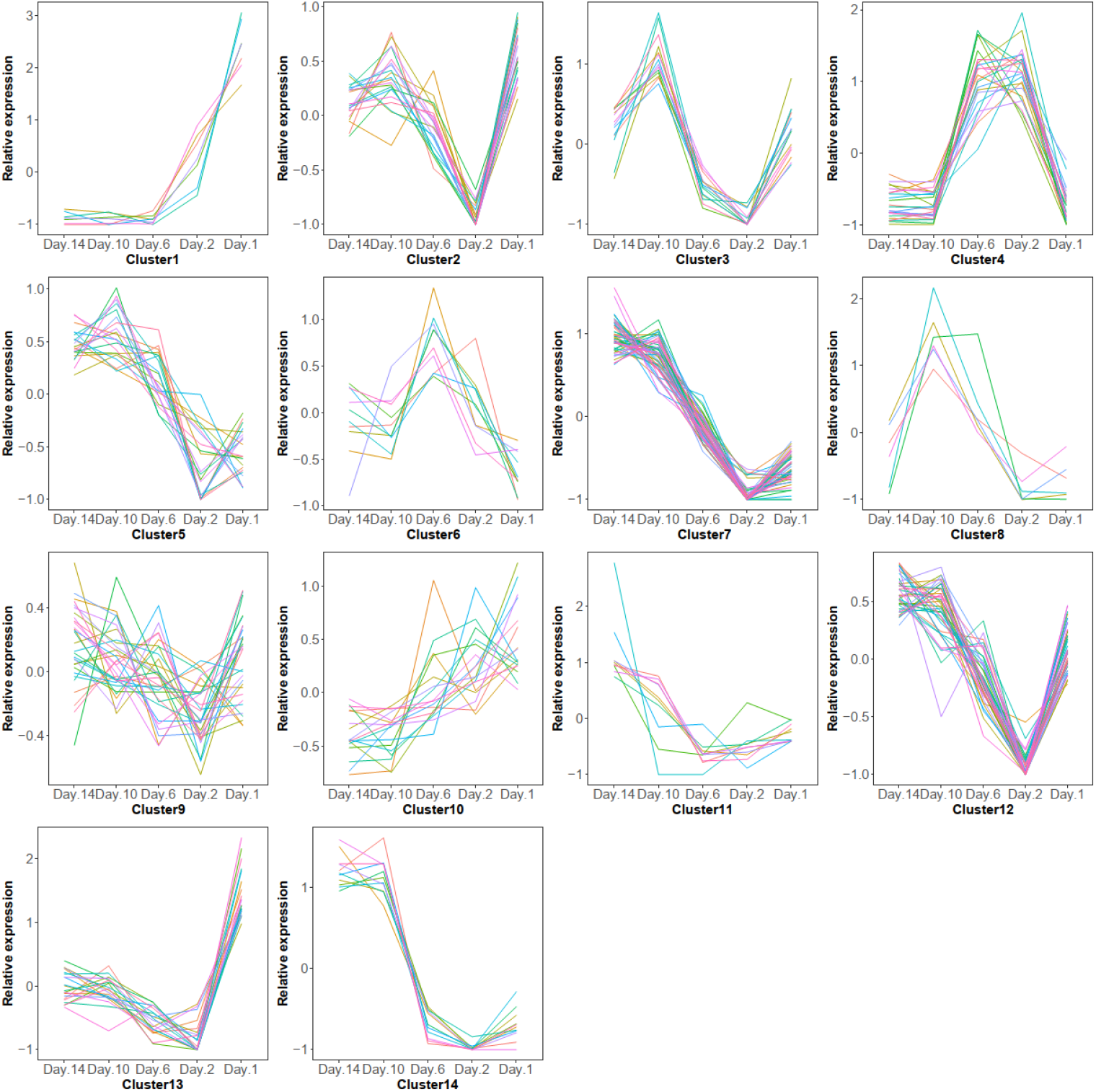
Expression pattern of differentially expreesed (DE) lncRNAs across five stages. The colored lines represent that the different lncRNA changes.

### Prediction of target genes

To investigate the probable roles of the lncRNAs in swine mammary gland, we predicted the target genes of lncRNAs. First, a search for protein-coding genes within 100 kb upstream/downstream of lncRNAs, and 6,589 genes were identified. Further, the expressions correlation between lncRNAs and protein-coding genes were examined, and 804 significant lncRNA-mRNA correlation pairs (*P* < 0.05) were detected, including 685 protein coding genes and 443 lncRNAs. Among the 443 lncRNAs, 149 were differentially expressed, including 5, 41, 94 and 109 DE lncRNAs respectively corresponded to −10vs-14, −6vs-14, −2vs-14 and +1vs-14 (Table S4).

In addition, we found that 256 protein-coding genes within 100 kb upstream/downstream of the 149 DE lncRNAs were also confirmed to be strongly correlated with them. To assess the function of DE lncRNAs, GO and KEGG enrichment analyses were performed using the 256 target genes with KOBAS. There were 311 significant enrichments were presented (*P* < 0.05; Table S5), including 298 GO terms and 13 KEGG pathways. These GO and KEGG enrichments were involved in 87 genes, and most of them were directly related to the labor and colostrum. For example, *CSN3, SMO*, and *CSN2* were involved in mammary associated activities.

### Module construction

To explore the specific lncRNAs or genes that were highly associated with lactation, we performed WGCNA using DE RNAs and their target genes. A total of 12 modules associated with the specific expression profiles of different samples were identified (Figure 4A). Then, we calculated associations of each module with five lactation stages, and found seven modules including Greenyellow, Green, Black, Yellow, Brown, Blue, and Turquoise, were significantly associated with Day −14, Day −2 and Day +1 (*P* < 0.05; Figure 4B). In detail, the Greenyellow module, including 12 lncRNAs and 6 target genes were highly associated with Day −2 (*P* = 0.02). The Green (including 5 lncRNAs and 30 target genes), Black (including 1 lncRNAs and 23 target genes), Yellow (including 5 lncRNAs and 49 target genes), and Brown (including 1 lncRNAs and 53 target genes) modules were significantly associated with the mammary gland samples at Day +1 (*P* =3e-04 ∼ 0.009). The Turquoise (including 9 lncRNAs and 91 target genes) and Blue (including 2 lncRNAs and 67 target genes) modules were strongly associated with at Day −14 and Day +1 (*P* = 0.004 ∼ 0.04). The lncRNAs and target genes involved in the significant modules were shown in Table S6.

**Figure 4.**
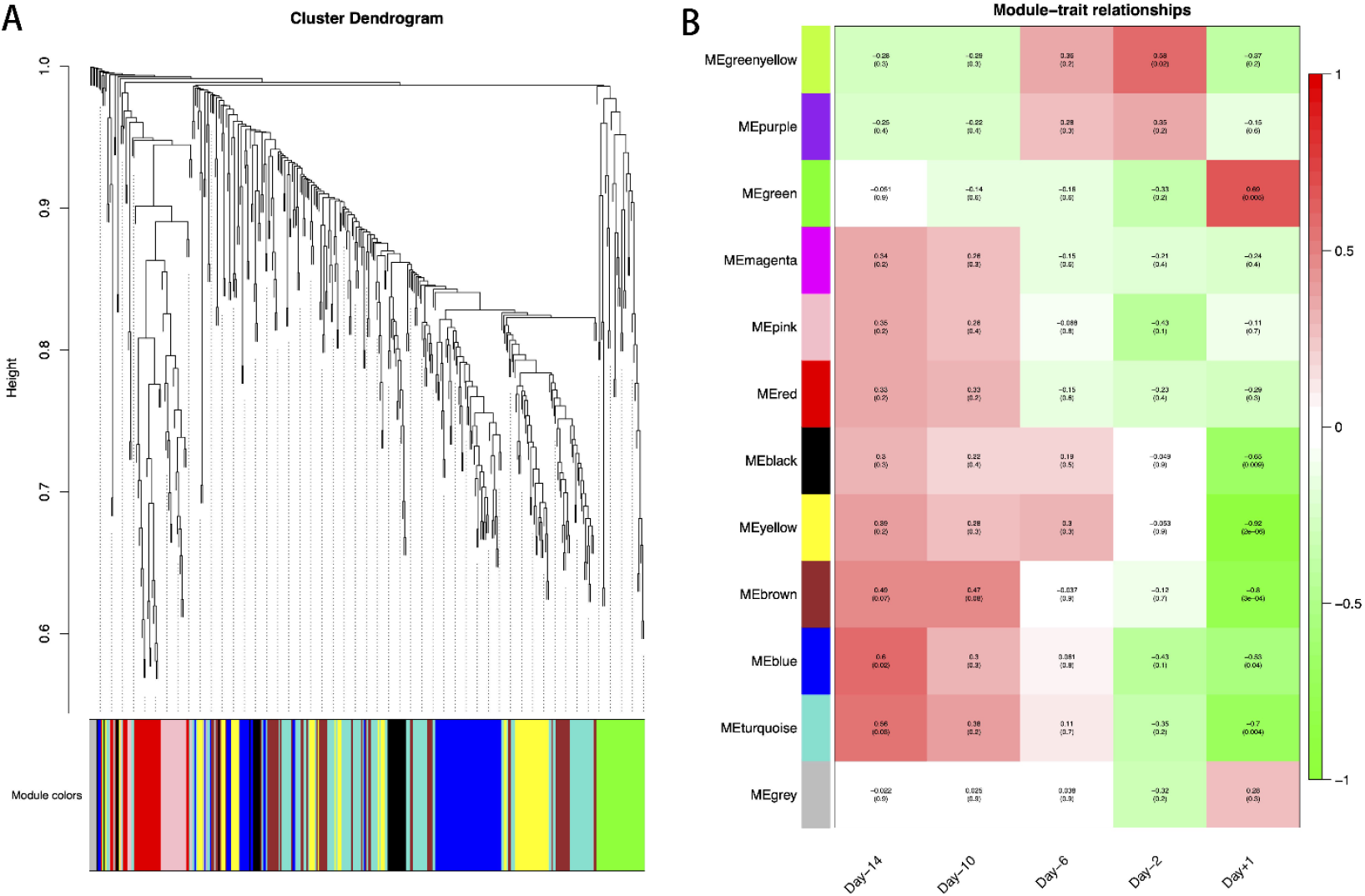
Weighted gene co-expression network analysis (WGCNA) at each swine mammary gland development. **(A)** Hierarchical cluster tree showing co-expression modules identified by WGCNA. **(B)** Module-sample relationships. Each column indicates a sample, and each row represents a module. The correlation coefficient and *p*-value of each sample-module relationship are displayed. Red represents high correlation value, and green represents low correlation value.

### Comprehensive analysis

Among the 256 target genes, 121 were differentially expressed across five lactation stages, and function annotation revealed that 25 of them targeted by 23 lncRNAs were involved in lactation developing, milk fat metabolism and colostrum function (Table 1). Interestingly, 20 target genes (*ACTN4, ADCY1, CSN3, SMO, PTK7, MPDZ, NPR1, CSN2, ATP2C2, PRKAG2, CD36, NPR1, ACSL3, GALNT15, C6H1orf210, CDC20, SNRPD1, FIBCD1, ACTB* and *GALNT7*) and 3 lncRNAs (*XLOC_025150, ENSSSCG00000011196*, and *XLOC_010589*) were also belong to significant modules by WGCNA. These 20 target genes were correspond to 18 lncRNAs, which also included the 3 lncRNAs (*XLOC_025150, ENSSSCG00000011196*, and *XLOC_010589*) involved in significant modules. Hence, we suggested that these 18 lncRNAs targeting 20 genes were the candidates involved in lactation of sow, which were considered as the potentially functional candidates impacting the mammary gland developments from colostrogenesis to lactogenesis. In addition, we conducted the network of promising candidate lncRNAs, genes and pathways (Figure 5), in which, the functions of lncRNAs were clearly shown. For example, XLOC_025150 might regulate *CSN3* gene, were involved in the mammary gland development and lactation.

**Table 1.**
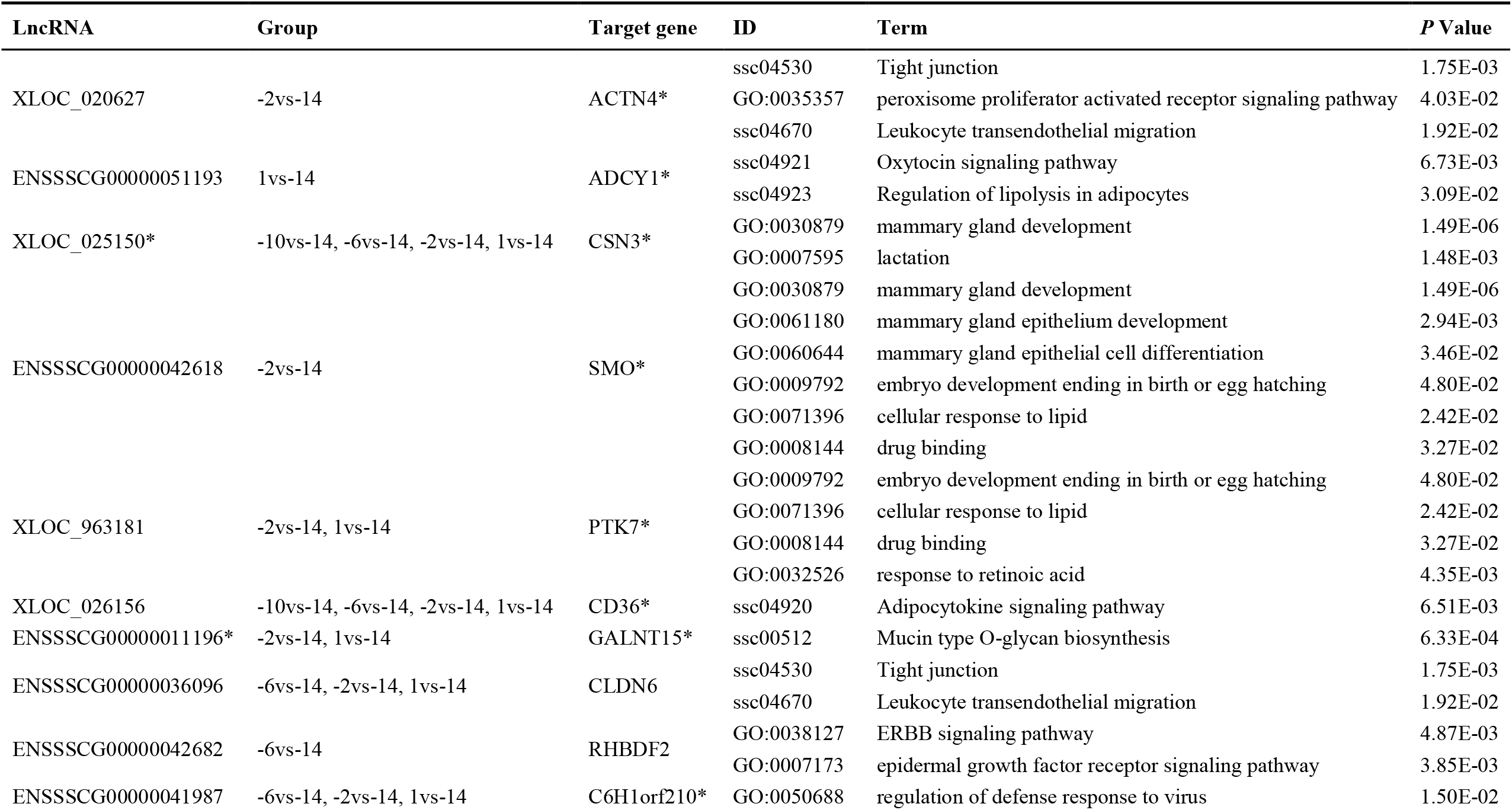
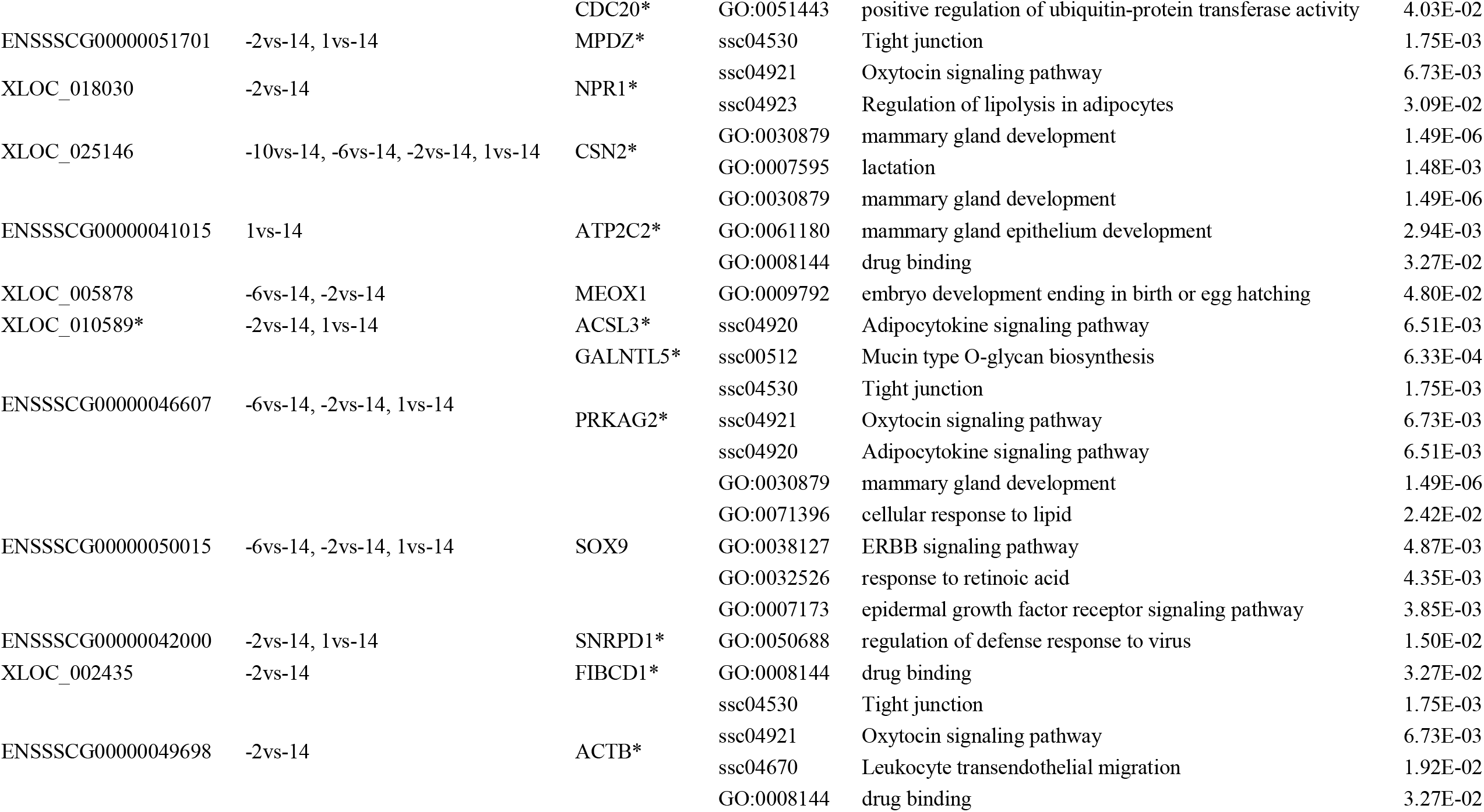
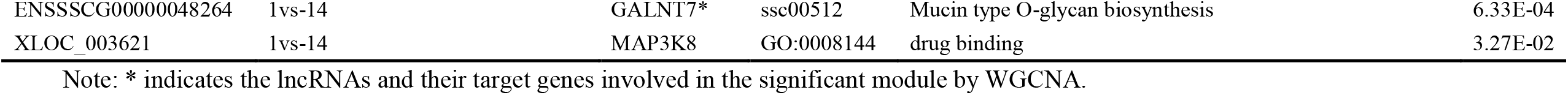
GO and KEGG enrichments of 23 potentially functional lncRNAs and their 25 target genes

**Figure 5.**
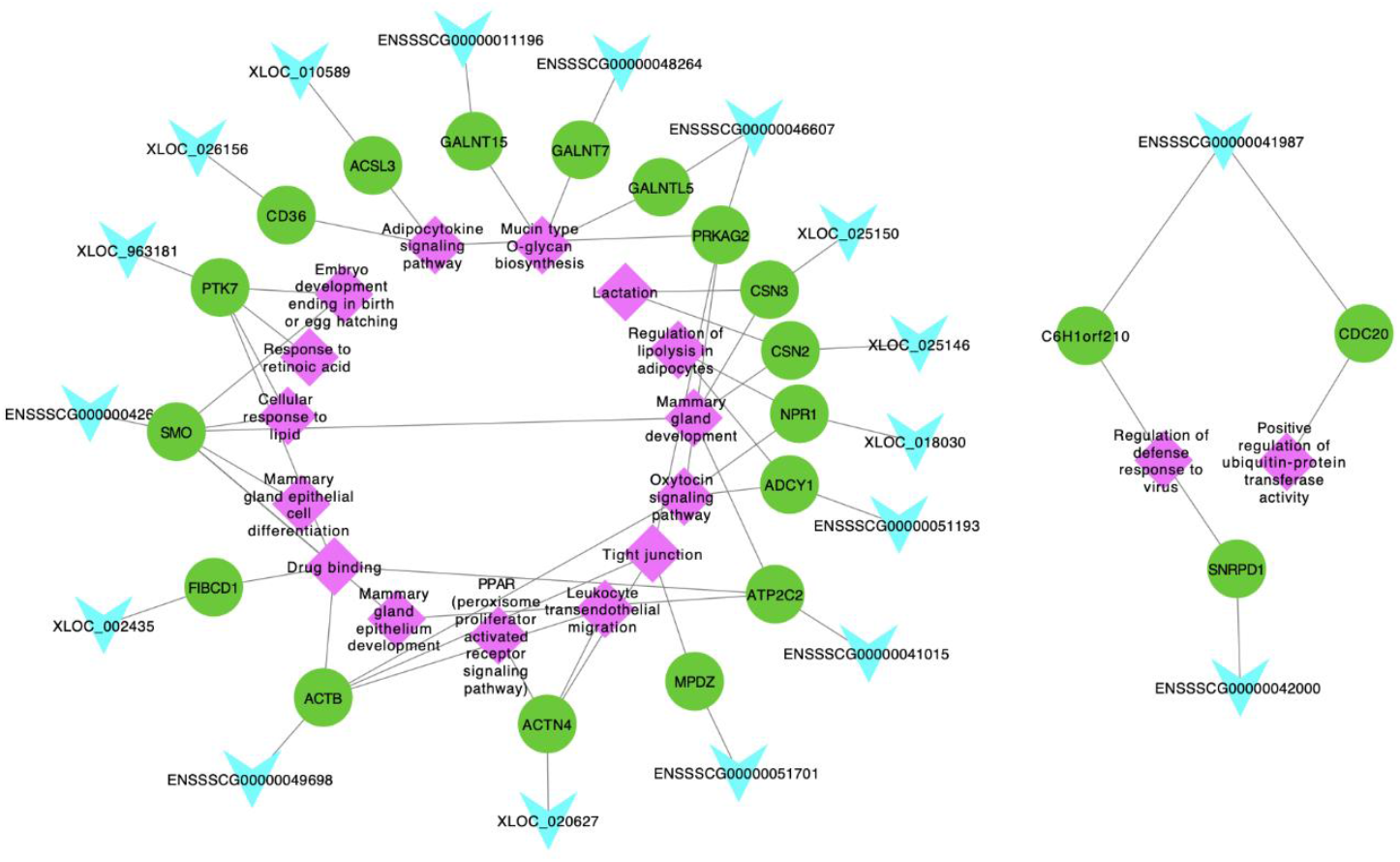
Network plot of the promising candidate lncRNAs, genes and pathways. The light blue, green and purple indicate lncRNAs, genes and pathways, respectively.

## Discussion

In this study, we systematically analyzed the RNA-seq data of 15 swine mammary gland samples collected at days 14, 10, 6 and 2 before (-) parturition to day 1 after (+) parturition, and identified 286 DE lncRNAs targeting 256 genes. Further, we found that these lncRNAs were involved in delivery and lactation developing, milk lipid metabolism, and immune function of colostrum, and 18 lncRNAs targeting 20 genes were proposed to be the candidates involved in lactation of sow.

At the parturition, the preparation for copious milk synthesis and secretion have begun (Kensinger et al. 1982; Palombo *et al*. 2018), and the mammary gland can reach the greatest degree of structural development. In this study, we examined the expression pattern of DE lncRNAs, most of the lncRNAs expressions were strongly changed from Day −2 to Day +1. Further, 18 promising functional lncRNAs targeting 20 genes were mainly identified in −2vs-14 and 1vs-14 groups, which reflected a strong activation of many metabolic processes before and after parturition. In the WGCNA, 20 promising functional DE target genes were involved in the significant modules, which were highly associated with the mammary gland samples at Day −14, Day −2 and Day +1. Hence, our results were consistent with the concept: before and after farrowing with the formation of colostrum, mammary glands undergo strong functional differentiation immediately (Hurley 2019).

In general, lncRNAs exert regulatory functions at different levels of gene expression to influence the tumor growth, cell-cycle, and apoptosis (Le *et al*. 2019; Wang et al. 2019b). For example, lncRNA H19 is involved in the regulation of high glucose-induced apoptosis through targeting VDAC1 (Li et al. 2016). Here, 11 lncRNA-mRNA target pairs, *XLOC_020627*-*ACTN4, ENSSSCG00000051193*-*ADCY1, XLOC_025150*-*CSN3, ENSSSCG00000042618*-*SMO, XLOC_963181*-*PTK7, ENSSSCG00000051701*-*MPDZ, XLOC_018030*-*NPR1, XLOC_025146*-*CSN2, ENSSSCG00000041015*-*ATP2C2, ENSSSCG00000046607*-*PRKAG2*, and *ENSSSCG00000049698*-*ACTB*, were involved in delivery and lactation metabolism, such as tight junctions, oxytocin, development of the mammary gland and lactation. Tight junctions of mammary gland from the pregnant animal are leaky, undergoing closure around parturition to become the impermeable tight junctions of the lactating animal (Nguyen & Neville 1998). In dairy cattle, after parturition, the start of copious milk production requires the closure of tight junctions to form the blood-milk barrier and prevent paracellular transfer of blood constituents into milk (such as lactate dehydrogenase and serum albumin) and vice versa (such as appearance of α-lactalbumin in blood) (Kessler et al. 2019). Oxytocin is a nonapeptide hormone that has a central role in the regulation of parturition and lactation (Arrowsmith & Wray 2014), and its best-known and most well-established roles are stimulation of uterine contractions during parturition and milk release during lactation (Gimpl & Fahrenholz 2001). Development of the mammary gland occurs in defined stages, which are connected to sexual development and reproduction (embryonic, prepubertal, pubertal, pregnancy, lactation, and involution) (Hennighausen & Robinson 2001). Lactation is a highly demanding lipid synthesis and transport process that is crucial for the development of newborn mammals (Wan et al. 2007). Embryo development ending in birth or egg hatching term was defined as the process whose specific outcome is the progression of an embryo over time, from zygote formation until the end of the embryonic life stage, and for mammals it is usually considered to be birth. The above reports suggested that these GO and KEGG enrichments were related with the parturition and lactation. Hence, we proposed that these lncRNAs in 11 lncRNA-mRNA target pairs might be regulated their target genes to impact the delivery and lactation processes.

Additionally, we found eight lncRNA-mRNA target pairs, *XLOC_020627*-*ACTN4, ENSSSCG00000051193*-*ADCY1, ENSSSCG00000042618*-*SMO, XLOC_963181*-*PTK7, XLOC_026156*-*CD36, XLOC_018030*-*NPR1, XLOC_010589*-*ACSL3*, and *ENSSSCG00000046607*-*PRKAG2*, were enriched in PPARs, regulation of lipolysis in adipocytes, cellular response to lipid, and adipocytokine signaling pathways, that might be related with the milk components metabolism.

For the piglets, colostrum plays an important role in ensuring their survival, growth and health by providing energy, nutrients, immunoglobulins, growth factors and many other bioactive components and cells (Quesnel & Farmer 2019). In the present study, we found that 12 lncRNA-mRNA target pairs, *XLOC_020627*-*ACTN4, ENSSSCG00000042618*-*SMO, XLOC_963181*-*PTK7, ENSSSCG00000011196*-*GALNT15, ENSSSCG00000041987*-*C6H1orf210, ENSSSCG00000041987*-*CDC20, ENSSSCG00000041015*-*ATP2C2, ENSSSCG00000046607*-*GALNTL5, ENSSSCG00000042000*-*SNRPD1, XLOC_002435-FIBCD1, ENSSSCG00000049698*-*ACTB* and *ENSSSCG00000048264*-*GALNT7*, were involved in leukocyte transendothelial migration, response to retinoic acid, drug binding, mucin type O-glycan biosynthesis, regulation of defense response to virus, and positive regulation of ubiquitin-protein transferase activity enrichments, which might impact the immune function of colostrum.

Based on these advantages, we proposed these 18 lncRNAs (*XLOC_020627, ENSSSCG00000051193, XLOC_025150, ENSSSCG00000042618, XLOC_963181, ENSSSCG00000051701, XLOC_018030, XLOC_025146, ENSSSCG00000041015, ENSSSCG00000046607, ENSSSCG00000049698, XLOC_026156, XLOC_010589, ENSSSCG00000011196, ENSSSCG00000041987, ENSSSCG00000042000, XLOC_002435*, and *ENSSSCG00000048264*) as the promising candidates for swine parturition and lactation. Our network results of 18 lncRNAs, their 20 target genes and corresponding pathways further clearly showed the functions of the candidate lcnRNAs and their target genes. There is a high probability that these lncRNAs and genes interact closely and thus act as key regulators in mammary gland, thus, future work should focus on the potential roles of them in sow lactation.

## Conclusions

Sow milk production is the major factor which can limit growth and survival of piglets. To enhance milk performance, it is necessary to understand the process of mammary gland morphogenesis and to identify the regulatory factors of mammary development. In this study, we identified 1,084 lncRNAs in swine mammary gland using RNA-seq data, and 286 were DE lncRNAs in five lactation stages. Integrated analysis of the DE lncRNAs expression pattern examination, targets prediction, function annotation and WGCNA, we proposed that 18 lncRNAs (such as *XLOC_020627, ENSSSCG00000051193, XLOC_025150, ENSSSCG00000042618, XLOC_963181, ENSSSCG00000051701, XLOC_018030* and *XLOC_025146*) targeting 20 genes (such as *ACTN4, ADCY1, CSN3, SMO, CSN2, PRKAG2, FIBCD1* and *GALNT7*) as the promising candidates were involved in swine parturition and colostrum occurrence processes. These results provided a new insight for exploring critically regulatory factors involved in reproductive performance of sow.

## Supplementary information

**Table S1** The information of RNA sequencing data for 15 samples of sow mammary gland tissue

**Table S2** The results of the identified lncRNAs in this study

**Table S3** The detailed information of DE lncRNAs in this study

**Table S4** The predicted target genes of DE lncRNAs

**Table S5** Significant GO and KEGG pathways of the target genes of DE lncRNAs

**Table S6** The lncRNAs and genes involved in significant modules

## Acknowledgements

This research was supported by the Agricultural Science and Technology Innovation Project (ASTIP-IAS02), and Chinese Academy of Agricultural Sciences Foundation (20118/2020-YWF-YTS-8).

## Data availability and materials

The data (BioProject ID: GSE101983) was in the Gene Expression Omnibus database.

